# WildTrust-UAV: uncertainty-aware wildlife detection with privacy-preserving spatial disclosure and verifiable inference provenance

**DOI:** 10.64898/2026.11.12.766079

**Authors:** Poonam Verma, Ravindra Nath Tripathi, Johnson Kinyua

## Abstract

Unmanned aerial vehicle (UAV) imagery has made wildlife surveys faster and safer, but a detected animal is not yet a conservation record. Detector confidence scores are often poorly calibrated, exact coordinates of threatened species can expose them to harm, and an observation is rarely tied back to the image, model, and settings that produced it. WildTrust-UAV addresses these three problems in one framework. It pairs a YOLO11 detector with stochastic-view uncertainty estimation, temperature scaling fitted on validation data, a selective gateway that accepts, defers, or rejects each detection, role-based disclosure of animal locations, and a provenance record that hashes the source image, model, configuration, prediction, uncertainty, timestamp, and nonce. Records are aggregated in a Merkle tree and signed with Ed25519 so that any observation can be verified independently. We report three evaluations on the public WAID dataset. In the detection experiment, two YOLO11n runs reached mAP@0.50 of 0.901 and 0.903 and mAP@0.50:0.95 of 0.527 on a 2,000-image held-out test set. In a separate reliability experiment on 122,720 test detections, temperature scaling reduced expected calibration error from 0.0715 to 0.0179 and the Brier score from 0.0353 to 0.0222. Composite uncertainty identified erroneous detections with an AUROC of 0.904, although confidence alone did better (0.992). Deferring the least reliable detections lowered selective risk from 0.722 at near-full coverage to 0.610 at 69.9% coverage, so considerable error remains in the candidate set. In the provenance experiment, the verifier rejected all 4,500 tampered submissions across nine attack scenarios and accepted all 500 genuine controls, with batch construction and signing taking 0.1 to 2.5 ms. Location policies used synthetic coordinates, and the ledger is a local prototype. Calibration and verifiable records can thus be added to a UAV detection pipeline at low cost, while combining uncertainty signals does not automatically improve error detection.

## Introduction

Unmanned aerial vehicles (UAVs) have become a common source of imagery in environmental remote sensing because they collect high-resolution images over large, remote, or hazardous terrain at modest cost. They can be flown repeatedly and below the altitude of conventional aircraft, which has opened new possibilities in mapping, habitat analysis, and wildlife population monitoring [1]. In ecological surveys, UAV imagery removes the need for observers to enter difficult landscapes and leaves a permanent photographic record that others can check. Controlled experiments have shown that UAV-assisted counts can be more accurate and more precise than traditional counting methods, particularly for animals spread over wide areas [2]. UAVs are therefore being used for population estimates, reserve monitoring, and the detection of poaching or trespass.

Collecting aerial imagery does not by itself produce useful monitoring data. A survey may generate thousands of high-resolution images in which animals occupy a tiny fraction of the frame, and their visibility depends on flight altitude, vegetation, shadow, background similarity, motion blur, illumination, animal density, and partial occlusion. Searching such images by hand is slow and prone to error and bias. Machine learning detectors are an obvious alternative, but wildlife imagery differs substantially from the datasets on which most computer vision methods are developed. Animals appear as small, scattered objects, and many features of the environment resemble them closely enough to produce false positives. Kellenberger et al. showed that the class imbalance and rarity of animals in UAV imagery require special care in both training and evaluation, beyond headline accuracy [3].

Benchmarks built for particular applications have improved aerial wildlife analysis. BIRDSAI provided annotated aerial thermal infrared video of humans and animals in African protected areas, supporting detection and tracking of individuals at night [4]. WAID collected 14,375 multi-class aerial drone images of sheep, cattle, seals, camels, kiang, and zebras for evaluating small-target wildlife detectors [5]. Kumar et al. developed WildlifeMapper, a flexible model for multi-species detection and identification across environments, demonstrating the value of models that transfer across regions and taxa [6]. Together these studies establish that automatic animal detection is feasible while identifying persistent difficulties: small targets, variable conditions, transfer across domains, and the gap between benchmark performance and operational conservation tasks. Most wildlife detection studies report precision, recall, and mean average precision. These metrics are necessary, but they do not say whether the confidence attached to an individual detection reflects the probability that it is correct. Deep networks can be discriminatively accurate yet overconfident [7]. Overconfidence matters in wildlife monitoring because a detector may attach a high score to a wrong species label, and a missed or misreported observation of an endangered species can carry real ecological cost. Confidence calibration is therefore needed to separate accuracy from reliability.

Selective prediction offers a second line of defense. A system need not commit to every observation; it can accept predictions that are reliable enough and defer the doubtful ones to a wildlife expert. This creates a measurable trade-off between risk and coverage: fewer observations are accepted automatically, but those that are accepted should be more reliable [8]. The arrangement suits conservation workflows, where automated analysis is meant to complement specialist review rather than replace it. A trustworthy aerial monitoring system should therefore support three outcomes for each detection: automatic acceptance, expert review, and rejection.

Detection accuracy is only one property of a trustworthy system. UAV images may reveal nest sites, migration routes, water points, breeding areas, or the locations of endangered individuals. Unrestricted access to such information creates ecological and security risks. A system that protects imagery from alteration but releases coordinates freely offers no privacy to conservation work. Access to geographic detail should depend on the user’s role and purpose: a public user might see a regional location, a researcher a protected grid cell, and an authorized wildlife officer the exact coordinate when the task requires it. This means the privacy mechanism must sit inside the inference workflow rather than beside it.

Provenance is a further problem. Once an image has been processed, the resulting observation can be copied, edited, relabelled, or separated from the model and configuration that produced it. Ordinary database logs can record that an inference request occurred, but they cannot establish a cryptographically verifiable link between the original image, the trained model, the inference configuration, the confidence value, and the output. Blockchains have been examined as one way of attesting the integrity of remote-sensing images [9], using distributed ledgers and hash functions. Storing only an image hash, however, does not prove AI provenance, because the same image processed with a different configuration or preprocessing can yield a different result. Large aerial images and prediction artefacts cannot be written directly to a blockchain. Content-addressable storage such as IPFS refers to objects by the cryptographic identifier of their content and organizes them as Merkle directed acyclic graphs [10]. The artefact can then stay off-chain while only its hash appears on the ledger. Writing each observation as a separate on-chain record would still be wasteful in storage and transmission cost. A hierarchical approach based on Merkle trees avoids this: many commitments are held at the leaves and a single aggregated root represents the whole batch [11]. The structure allows an individual record to be verified with a number of hashes that grows logarithmically with the batch size.

A valid observation must also be attributable to an authorized sensor or processor. A digital signature binds the root of the evidence to its origin and allows independent verification without revealing the signing key. Ed25519, specified in RFC 8032, is a compact signature scheme on an Edwards curve with fixed parameters and published test vectors [12]. With fresh nonces and timestamps, signed evidence records can also expose replays. Together these mechanisms distinguish a genuine observation from one whose image was replaced, whose model was switched, whose prediction or configuration was edited, or whose earlier evidence record was reused as if new.

Although detection, uncertainty estimation, data storage, and cryptography have each advanced, they are usually treated as separate problems. Wildlife detection research concentrates on accuracy, while secure UAV architectures concentrate on communication, access logging, or distributed image storage. A sound conservation pipeline has to answer three questions at once: was an animal reliably detected, was sensitive location information released appropriately, and can the observation be attributed to a specific image, model, and configuration? Answering only one of them is not enough for practical use.

This paper presents WildTrust-UAV, a wildlife detection framework that combines uncertainty estimation, cryptographic evidence integrity, and role-based release of animal locations. The framework uses a multi-scale detector with optional overlapping tiled inference, calibrates confidence values, and applies a reliability gate that assigns each observation to automatic acceptance, expert review, or rejection. A role-based spatial policy restricts the precision at which animal locations are disclosed. For traceability, the framework builds an evidence commitment by hashing the source image, model parameters, inference configuration, predicted object, uncertainty value, timestamp, and a fresh nonce. Commitments are aggregated in a Merkle tree and signed with Ed25519. Large artefacts stay in off-chain content-addressable storage, and compact provenance data are anchored to a permissioned ledger.

The contributions of this work are:

- an uncertainty-aware wildlife monitoring design that integrates stochastic-view analysis, calibration, selective referral, and optional tiled inference, in which the components with completed comparative evidence are distinguished from those without;
- a spatial disclosure scheme that releases animal positions at a granularity set by user role and usage context;
- an evidence record that covers the source image, model version, inference setup, prediction, uncertainty score, timestamp, and nonce;
- Ed25519-signed Merkle batching for compact provenance verification, evaluated separately from detector correctness and confidence calibration; and
- a reproducible evaluation protocol on the WAID benchmark covering repeated randomized runs, calibration testing, risk-coverage analysis, computational profiling, component comparison, location-policy checks, and adversarial attacks on provenance.

Unlike a blockchain store for images, WildTrust-UAV treats detection reliability, location privacy, and inference provenance as related problems. The aim is not to argue that blockchain or object detection alone makes conservation decisions trustworthy, but to propose a measurable architecture in which each property can be assessed on its own and in combination.

### Related work

#### UAV-based wildlife monitoring

UAVs give ecologists an efficient way to photograph large or inaccessible habitats at high resolution while reducing the time and risk of traditional surveys. With machine learning, these image collections can be used to identify species, estimate abundance, and map spatial distributions [13]. Ecological images differ sharply from standard object detection benchmarks: animals may be only a few pixels wide, blend into vegetation, occur in highly unbalanced ratios, and be partly hidden by canopy or terrain.

Eikelboom et al. showed that deep learning applied to aerial images produces population estimates that are more precise and more accurate than manual analysis [14]. Kellenberger et al. found that active learning can cut the labelling effort for animal detection in UAV imagery, with a small set of strategically chosen labels enough to train an effective detector [15]. The AIDE platform provides a practical framework for training classifiers and detectors for ecological surveys and shows the value of human-in-the-loop model development [16]. None of these studies combined accurate prediction with uncertainty estimates, privacy-preserving spatial information, and per-prediction provenance.

WAID is one of the first benchmarks for wildlife detection in aerial images, with several animal categories and realistic variation in scale, background, and density [5]. High performance on a fixed test split does not, however, guarantee reliability. Images captured at different altitudes, seasons, sensors, locations, or habitats may come from a different distribution than the training set, and a detector with high mean average precision can still be confidently wrong in deployment.

#### Small-object detection in complex habitats

Small-object detection remains a central technical difficulty in aerial wildlife monitoring. Weak spatial features of distant animals can be lost to downsampling in convolutional backbones, while rocks, shrubs, shadows, and reflections on water can resemble animals. Multi-scale feature fusion, high-resolution feature maps, image tiling, and context-aware augmentation are widely used to preserve small-object information [3,5].

One-stage detectors offer a good balance between accuracy and computational cost, but their standard training objectives are not aligned with ecological risk. Classification and localization losses penalize individual object-level errors, whereas in practice a missed animal, a duplicate detection, a species confusion, and a false alarm in a high-value area have very different consequences. Aggregate mean average precision also hides differences between common and rare species; a detector can have excellent mAP and still perform poorly on the rare classes that matter most.

WildTrust-UAV therefore keeps an efficient multi-scale detector as its visual core and adds three components aimed at deployment: uncertainty estimation, risk-aware selection, and verifiable inference recording. The design does not assume that more architectural complexity yields better detection. The detector is one part of a larger decision-support system in which predictions are calibrated, selectively disclosed, protected geographically, and verified.

#### Predictive uncertainty and selective inference

Confidence values from modern neural networks do not necessarily match empirical correctness rates. Guo et al. showed that high-performing classifiers can be poorly calibrated and that post hoc temperature scaling can correct this without changing the predicted classes [7]. The point matters for wildlife monitoring because confidence may decide whether an observation is accepted automatically, sent to an expert, or discarded.

Deep ensembles estimate epistemic uncertainty by combining predictions from independently trained models [17]. They improve calibration and robustness, but their computational and storage demands are unsuited to UAV or edge deployment. A cheaper option is to generate stochastic predictions through test-time augmentation or dropout and to measure disagreement across inferences. These methods do not provide a true posterior, but they do give useful information about prediction stability.

Selective prediction allows a model to abstain when the estimated risk of error is too high. SelectiveNet builds a rejection function into the network and minimizes risk under a target coverage constraint [18]. In wildlife applications, abstaining is preferable to forcing a wrong species label, because the resulting observations feed population estimates and other conservation decisions. Such models are evaluated with risk-coverage curves: as a larger share of detections is accepted, the error rate among accepted predictions rises.

Calibration and selective prediction have mostly been studied for image-level classification. Detection adds further sources of uncertainty: class probabilities, bounding-box localization, objectness, non-maximum suppression, and disagreement between overlapping predictions. WildTrust-UAV addresses this with a detection-level uncertainty score built from calibrated class probability, localization stability under stochastic transformations, and cross-view agreement. Observations whose reliability falls below the acceptance threshold are sent to an expert review queue rather than dropped.

#### Privacy-aware disclosure of wildlife locations

Exact wildlife locations can be sensitive. Publishing breeding sites, migration routes, nests, or the positions of endangered animals raises the risk of disturbance, poaching, unauthorized sample collection, and habitat exploitation. Biodiversity data guidelines therefore advise against disclosing precise coordinates when disclosure could harm the species recorded [19]. Stripping all geographic metadata is not a solution for collaborative monitoring, because authorized conservation officers need exact coordinates to verify observations and act on them. Leaving exact coordinates open in a shared detection report creates risk in the other direction. A suitable architecture distinguishes three views of the same observation: a public view, a researcher view, and an operational view for authorized staff.

WildTrust-UAV uses a hierarchical spatial disclosure policy. Exact coordinates are available only to authorized users; everyone else receives a generalized location such as a grid cell, a geohash prefix, or a deliberately perturbed coordinate. The disclosure level depends on the sensitivity category of the species and the recipient’s authorization. The framework also keeps a cryptographic commitment to the original location, so that a disclosed or decrypted coordinate can be checked against the initial record.

#### Blockchain and verifiable data provenance

Blockchains have been explored for UAV coordination, surveillance, identity management, and secure communication [20]. De Campos et al. described a blockchain-enabled multi-UAV surveillance system in which autonomous UAVs coordinate through smart contracts [20]. Such work demonstrates distributed coordination but does not address whether an individual wildlife prediction was produced by a specific model from an untampered input.

Data-provenance frameworks rely on cryptographic hashing and append-only logs to detect alteration. ProvChain, for example, uses blockchain to provide accountability and provenance validation for cloud data [21]. Provenance models more generally represent the relationships among entities, the computational activities that produced them, and the agents responsible [22]. The same ideas apply to ecological AI: a wildlife record should carry the input image, model version, inference parameters, timestamp, approximate acquisition location, and responsible agent or device.

Writing full UAV imagery and prediction files to a blockchain is impractical: the files are large, transaction costs can be high, and immutable publication conflicts with privacy needs. WildTrust-UAV therefore separates on-chain and off-chain data. Images, encrypted coordinates, uncertainty vectors, and detailed inference reports stay in controlled off-chain storage; only the cryptographic commitment and audit metadata go to the ledger. Many inference records are aggregated in a Merkle tree under a single root, and a verifier can check one record with its Merkle proof without seeing the others.

A blockchain does not prove that a detection is scientifically correct, nor that the sensor acquired the event authentically. Its role is narrower but still useful: tamper evidence, temporal ordering, traceability of model version, and non-repudiable publication. The validity of the detection must be assessed separately.

#### Research gap

Prior work has separately advanced UAV wildlife detection, active annotation, uncertainty quantification, selective prediction, spatial data confidentiality, and blockchain provenance. To our knowledge, no existing framework integrates these within a single ecological inference pipeline that can support population estimation and conservation decisions. Detection studies report accuracy without cryptographic proof of the model and data behind a given result, and blockchain-based UAV studies secure communication and coordination without analysing detector calibration, selective risk, or location sensitivity.

WildTrust-UAV closes this gap by combining (1) multi-scale wildlife detection; (2) uncertainty estimation and calibration at the level of individual detections; (3) risk-controlled acceptance, abstention, and referral of uncertain observations to experts; (4) spatial disclosure tailored to user role and species sensitivity; and (5) inference provenance secured by Merkle-batched, digitally signed records. The novelty lies in the experimental setup that evaluates accuracy, calibration, selective validity, privacy, overhead, and provenance together, rather than in blockchain-supported detection alone.

## Materials and methods

### Problem formulation

Let the wildlife detection dataset be

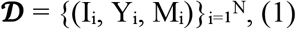

where Iᵢ is the i-th aerial image, Yᵢ its ground-truth annotations, and Mᵢ optional acquisition metadata such as timestamp, UAV identifier, altitude, camera parameters, and geographic coordinates. The annotations are

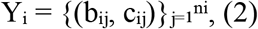

where bᵢⱼ ∈ ℝ⁴ is the bounding box of the j-th wildlife instance, cᵢⱼ ∈ {1, 2, …, C} its species class, nᵢ the number of annotated instances in image Iᵢ, and C the number of wildlife classes.

Given image Iᵢ, the detector fθ produces

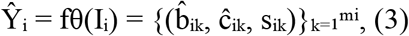

where b^ᵢₖ ∈ ℝ⁴ is a predicted bounding box, ĉᵢₖ ∈ {1, 2, …, C} the predicted class, sᵢₖ ∈ [0, 1] the raw confidence score, and mᵢ the number of candidate detections for image Iᵢ.

The raw score sᵢₖ in Eq (3) may not represent the true probability that the detection is correct, so a calibration function gφ: [0, 1] → [0, 1] is applied:

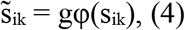

where s̃ᵢₖ is the calibrated confidence. A prediction is retained when its calibrated confidence exceeds an operating threshold τ:

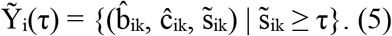

Conventional detectors return Ŷᵢ in Eq (3) as their final output. WildTrust-UAV instead produces an enriched observation for every candidate detection:

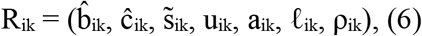

where uᵢₖ ≥ 0 is the composite uncertainty, aᵢₖ the selective decision, ℓᵢₖ the privacy-controlled location representation, and ρᵢₖ the cryptographic provenance commitment. The selective decision takes one of three values:

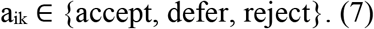

Thus, in addition to a box, class, and confidence, each observation carries its reliability, an operational decision, privacy-preserving location information, and verifiable provenance. The overall design objective can be written as

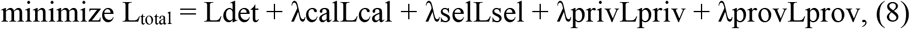

where Ldet is the wildlife detection loss, Lcal the calibration loss, Lsel the selective-risk loss, Lpriv the location-privacy loss, and Lprov the provenance-verification loss. The non-negative coefficients λcal, λsel, λpriv, and λprov weight the corresponding objectives. The framework aims to maximize detection performance while minimizing calibration error and selective risk and preserving location confidentiality and record verifiability.

### Overall system architecture

WildTrust-UAV has five modules: a multi-scale wildlife detection module; a stochastic-view uncertainty module; a calibration and selective decision gateway; a privacy-sensitive spatial disclosure module; and a verifiable inference-provenance module. Fig 1 shows the processing workflow.

**Fig 1.**
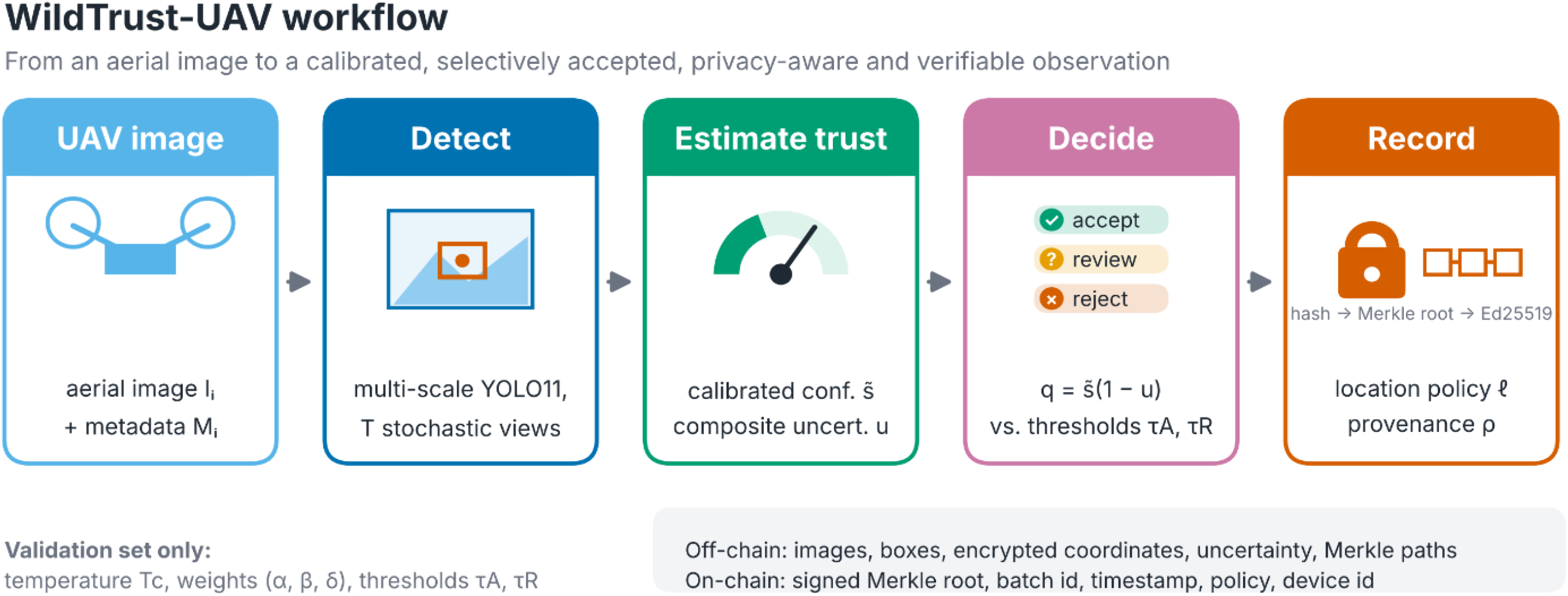
Processing workflow of WildTrust-UAV. An aerial image passes through the multi-scale detector, trust estimation (confidence calibration and composite uncertainty), the selective decision gateway, and the trustworthy record stage, which applies the spatial disclosure policy and produces the signed provenance commitment.

An aerial image is analysed by a multi-scale YOLO detector on the original view and on a set of transformed views. Matched detections across views are used to estimate classification, localization, and detection-stability uncertainty. Calibrated detections are then accepted, deferred to expert review, or rejected. Location information is released according to species sensitivity and user authorization, and a canonical inference record is hashed, batched in a Merkle tree, and digitally signed.

The uncertainty and provenance modules measure different things. The uncertainty module estimates how likely a prediction is to be correct; the provenance module checks whether the record has been altered. Validity and integrity are independent: a cryptographically valid record can contain a wrong detection, and an accurate prediction cannot be verified without its provenance. WildTrust-UAV reports the two properties separately.

### Multi-scale wildlife detector

YOLO11 is the detection backbone because it combines multi-scale feature extraction with fast one-stage inference. The network is pretrained and then fine-tuned on the WAID training subset. Its feature hierarchy preserves the high-resolution spatial detail needed to find small animals and the deeper semantic features needed to separate species.

### Detector training objective

For each training image the detector minimizes a composite loss:

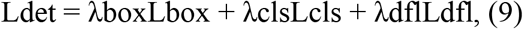

where Lbox penalizes localization error, Lcls is the classification loss, and Ldfl is the distribution focal loss used for box regression. The coefficients λbox, λcls, and λdfl set the relative weight of each term.

To preserve small wildlife instances during training, we use multi-scale variation, horizontal flipping, photometric transformations, and mosaic augmentation. Mosaic augmentation is applied only to the training set. All confidence thresholds, calibration parameters, uncertainty weights, and abstention thresholds are chosen on the validation set alone; the test set is used once, for the final evaluation.

### Stochastic-view uncertainty estimation

A single confidence score cannot fully describe the reliability of a detection. WildTrust-UAV therefore estimates uncertainty from T repeated inferences under a set of label-preserving image transformations:

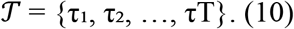

The set may include the identity, horizontal reflection, moderate brightness and contrast changes, and small rescaling. Transformations that could change the ecological interpretation of an image or erase very small animals are excluded.

The trained detector processes each transformed view:

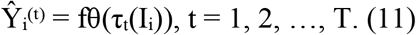

All predicted boxes are mapped back to the coordinate system of the original image. Detection k in the reference view is compared with candidate detection j in transformed view t; the two are matched when they share a predicted class and their intersection over union (IoU) is at least γ:

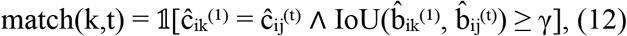

where 1[·] is the indicator function and γ the matching threshold. Each retained detection then receives three complementary uncertainty estimates.

### Classification uncertainty

Let pᵢₖ⁽ᵗ⁾ be the class-probability vector of detection k in view t, and Tᵢₖ the number of views in which the detection is matched. The mean class-probability vector is

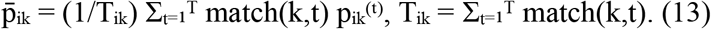

The normalized predictive entropy of the mean distribution is

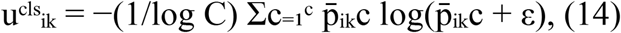

where ε is a small positive constant. Values near zero indicate a concentrated class distribution; values near one indicate strong class ambiguity.

When full class-probability vectors are not exposed by the detector interface, classification uncertainty is approximated by the variance of the matched confidence scores:

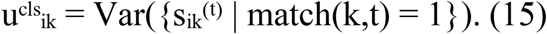

The reliability experiment reported here uses the confidence-variance approximation of Eq (15) rather than the entropy of Eq (14). Raw confidence variance is normalized with the validation 1st and 99th percentiles, u = clip((v − Q₀.₀₁,val)/(Q₀.₉₉,val − Q₀.₀₁,val + ε), 0, 1), and the same frozen bounds are applied to test detections. A raw variance of 1.0 is assigned when fewer than two matched views are available.

#### Localization uncertainty

Let

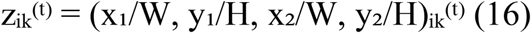

be the normalized corner coordinates of a matched box. Raw localization dispersion is the mean variance across the four coordinates:

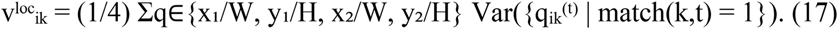

This raw dispersion is scaled with the validation 1st and 99th percentile bounds and clipped to [0, 1], as for Eq (15); test detections use the same bounds. Fewer than two matched views give a raw dispersion of 1.0. This is the implementation used to produce the reported results; it does not divide variance coordinate by coordinate.

#### Detection-instability uncertainty

A real animal should remain detectable under mild label-preserving transformations. Stochastic-view detection stability is

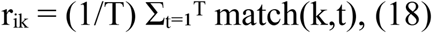

and the corresponding instability uncertainty is

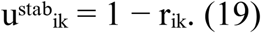

every view has uˢᵗᵃᵇᵢₖ = 0; one found only in the reference image receives a high value.

#### Composite uncertainty

The three components are combined as

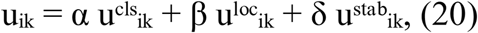

subject to

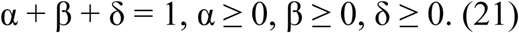

The weights are chosen on the validation set to maximize the ability of uᵢₖ to rank incorrect detections above correct ones. For this purpose, a prediction is correct when its class matches the ground-truth class and its IoU with the ground-truth box is at least 0.5:

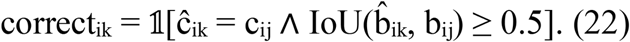

The stochastic-view estimator runs on a single trained detector and needs no ensemble retraining. Whether the extra forward passes improve error ranking over confidence alone is tested empirically in the Results; neither computational savings nor superior reliability is assumed.

### Confidence calibration

Raw detector confidence may be systematically over- or underconfident. WildTrust-UAV applies temperature scaling fitted on the validation set [7]. For detection logit zᵢₖ, calibrated confidence is

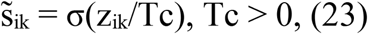

where σ(·) is the sigmoid function and Tc the learned temperature, obtained by minimizing thenegative log-likelihood over validation detections:

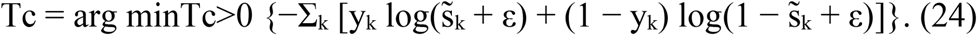

Here yₖ = 1 marks a correct detection and yₖ = 0 an incorrect or unmatched one.

Calibration quality is measured with the Brier score, negative log-likelihood (NLL), and expected calibration error (ECE). For Nd evaluated detections,

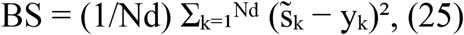

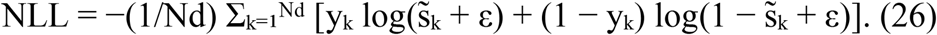

For ECE, detections are partitioned into B non-overlapping confidence bins S₁, …, SB with empirical accuracy and mean confidence

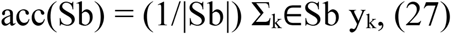

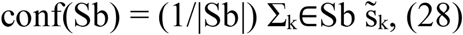

and

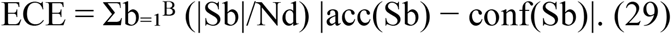

Lower values of Eqs (25), (26), and (29) indicate better calibration. All calibration parameters are learned from the validation set only; fitting Tc on the test set would leak information and is not done.

### Risk-aware selective decision gateway

Calibrated confidence and stochastic-view uncertainty are combined into a reliability score:

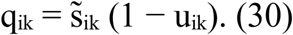

A detection scores high only when it has both high calibrated confidence and low composite uncertainty. Each detection is then assigned one of three decisions:

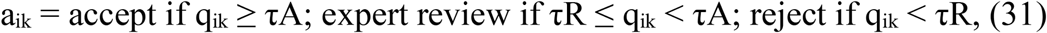

where τA and τR are the acceptance and rejection thresholds, 0 ≤ τR < τA ≤ 1, both chosen on validation data. Accepted observations enter automated ecological reports, deferred ones go to a human review queue, and rejected predictions are kept only in the audit record.

For an operating threshold τ, coverage is

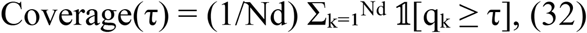

and selective risk is the share of incorrect predictions among accepted detections:

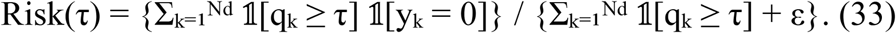

A good selective detector shows falling risk as coverage is reduced. WildTrust-UAV reports the full risk-coverage curve rather than a single hand-picked threshold. The area under the risk-coverage curve is

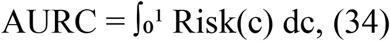

where c is coverage; a lower AURC means incorrect detections are assigned low reliability, and so deferred or rejected, before correct ones.

### Privacy-aware spatial disclosure

Each verified observation may carry an exact geographic coordinate:

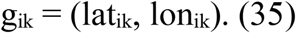

Every detected species is assigned a geographic sensitivity category from a configurable species-policy registry:

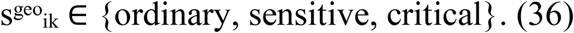

The disclosed location depends jointly on the exact coordinate, the species sensitivity, and the requesting user’s authorization:

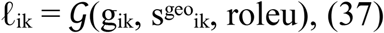

where G is the spatial-disclosure function and roleu the user’s role. For an ordinary species, an authorized research user may receive the exact coordinate. For a sensitive species, the coordinate is generalized to a grid cell or a truncated geohash. For a critical species, the public record contains only a coarse administrative region, and the exact coordinate remains encrypted and accessible only to authorized conservation staff.

For a chosen resolution r, grid-based generalization is

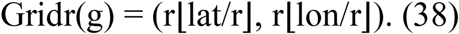

A smaller r gives finer spatial precision; a larger r gives stronger generalization. The exact coordinate is protected with authenticated encryption:

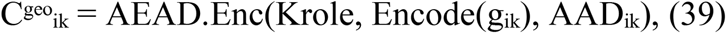

where Krole is the key assigned to the authorized role and AADᵢₖ is additional authenticated data binding the ciphertext to the observation identifier, device, and policy version. A salted geographic commitment is also generated:

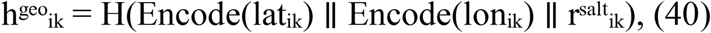

where H(·) is a cryptographic hash, ∥ concatenation, and rˢᵃˡᵗᵢₖ a high-entropy random salt. The salt matters because the set of plausible coordinates within a small region may be small enough to permit a dictionary attack on an unsalted hash.

### Verifiable inference provenance

For every observation, WildTrust-UAV builds a canonical provenance record:

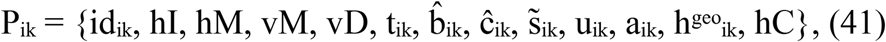

where idᵢₖ is the observation identifier, hI the input-image hash, hM the trained-model hash, vM the model version, vD the dataset version, tᵢₖ the inference timestamp, and hC the hash of the configuration used for preprocessing, inference, calibration, uncertainty estimation, and selective thresholding.

The record is serialized with canonical key ordering, explicit data types, standardized coordinate encoding, and deterministic numeric precision. Its leaf commitment is

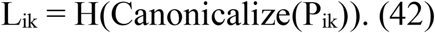

For a batch of n leaves, the level-zero Merkle nodes are

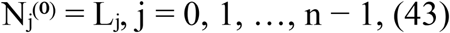

and adjacent nodes are combined recursively,

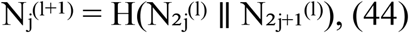

until a single root RB remains. When a level has an odd number of nodes, the last node is duplicated according to the declared tree-construction policy.

The UAV or an authorized ground station builds the batch descriptor

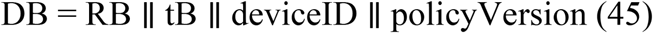

and signs it with its private key sk:

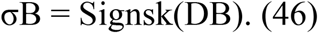

A verifier accepts an observation only when the batch signature is valid,

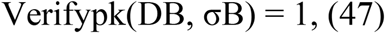

the leaf belongs to the committed tree,

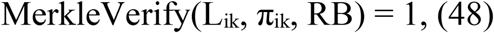

and all recomputed commitments agree with the record:

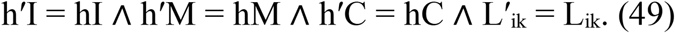

Here pk is the trusted public key and πᵢₖ the Merkle authentication path. Eqs (47) to (49) together verify batch authenticity, observation inclusion, and the integrity of the image, model, configuration, and record.

Only RB, the batch identifier, timestamp, policy version, device identifier, and signature σB need to be stored on the permissioned ledger. High-volume or sensitive material, including aerial images, bounding boxes, encrypted coordinates, full uncertainty records, and Merkle paths, stays off-chain. This gives verifiable provenance without exposing sensitive wildlife information or burdening the ledger.

### End-to-end inference flow

For an input aerial image, WildTrust-UAV proceeds as follows: compute the input-image hash; run YOLO11 inference on T stochastic views; project all detections back to the original coordinate system; match detections by class and IoU; compute classification, localization, and stability uncertainties; calibrate detection confidence; compute the reliability score and selective decision; apply the species- and role-specific spatial policy; compute the salted commitment to the exact coordinate; hash the canonical provenance record; include the record in a Merkle batch; and sign and commit the batch root. Fig 2 shows the full architecture.

**Fig 2.**
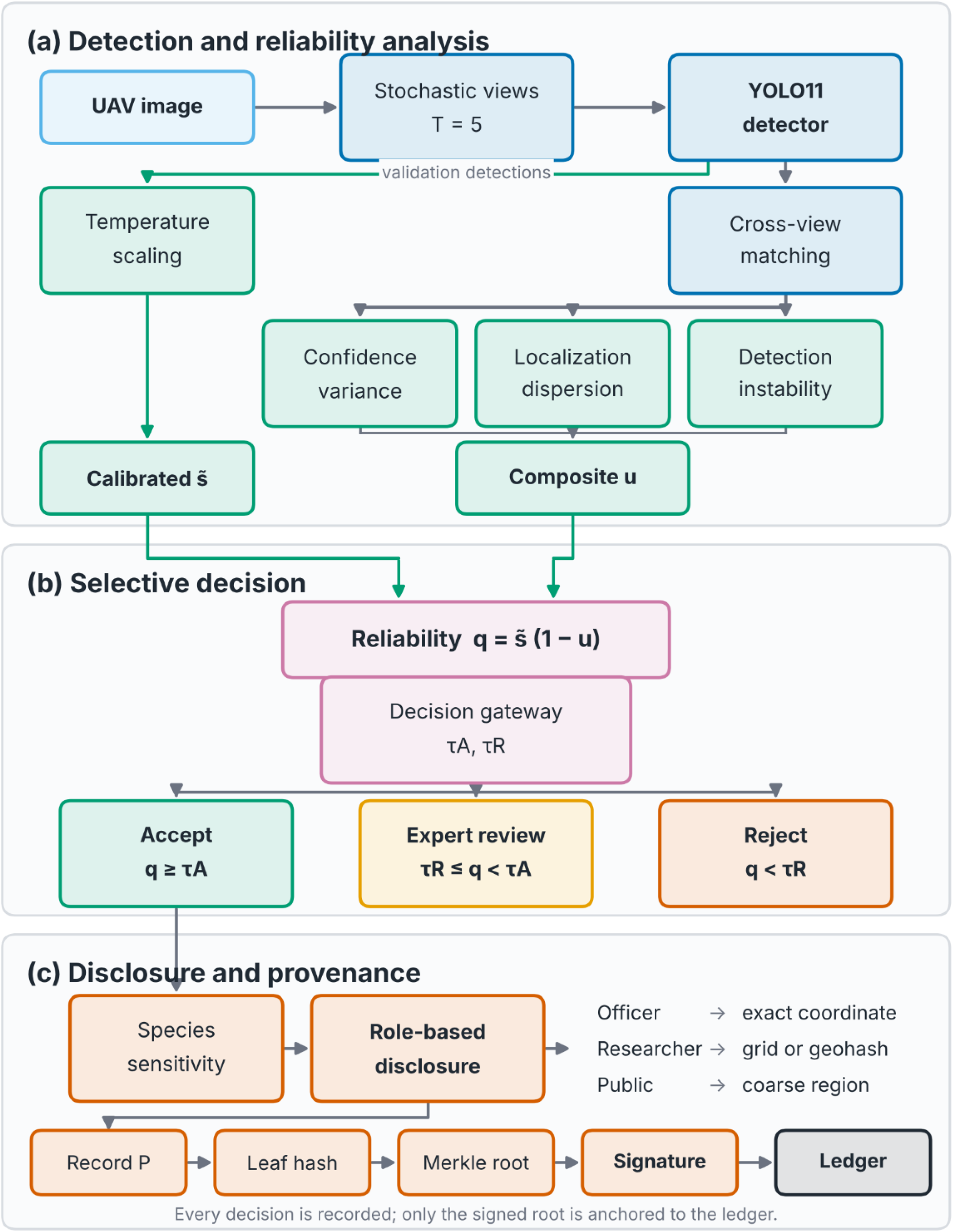
WildTrust-UAV architecture. (a) Detection and dual-source reliability analysis: stochastic views, multi-scale YOLO11 detection, cross-view matching, and the three uncertainty components. (b) Calibration, selective decision, and spatial disclosure: the reliability score q = s̃(1 − u) feeds the gateway that accepts, defers, or rejects each observation. (c) Privacy and verifiable inference provenance: role-based disclosure, canonical record construction, Merkle batching, signing, and anchoring to the permissioned ledger. Detailed images, uncertainty records, and encrypted coordinates remain in controlled off-chain storage; only the signed Merkle root and essential audit metadata reach the ledger.

The modular design lets each claim be tested on its own. Detection is measured by precision, recall, mAP@0.50, and mAP@0.50:0.95; reliability by calibration and risk-coverage analysis; privacy by disclosure and decryption tests; and provenance by tampering, signing, and Merkle-proof experiments.

### Research questions

The experiments address seven questions.

RQ1: Does WildTrust-UAV improve wildlife detection over regular lightweight detectors?

RQ2: Does stochastic-view uncertainty separate accurate detections from inaccurate or unstable ones?

RQ3: Does validation-based calibration strengthen the relationship between confidence and accuracy?

RQ4: Can selective prediction reduce detection risk effectively?

RQ5: What does each uncertainty component (classification, localization, and detection stability) contribute?

RQ6: Can sensitive wildlife locations be protected while records remain recoverable and verifiable?

RQ7: What overhead do uncertainty estimation and provenance computation add?

### Dataset and partitioning

Experiments use a public 8,000-image subset of the Wildlife Aerial Images from Drone dataset (WAID) [5], with object-level bounding-box annotations and six animal categories. Table 1 summarizes the partitions.

**Table 1.** WAID dataset partitions.

| **Partition** | **Number of images** | **Usage** |
|---|---|---|
| Training | 4,000 | Detector optimization |
| Validation | 2,000 | Model monitoring and calibration source |
| Test | 2,000 | Held-out evaluation only |
| Total | 8,000 |  |

No test image is used for model selection, temperature estimation, uncertainty-weight tuning, or selective thresholding. Where several frames come from one continuous UAV video sequence, sequence-level leakage across partitions is checked using filenames, metadata, and perceptual hashes.

Descriptive statistics for each partition cover the number of images and objects, class-wise object counts, class imbalance ratios, the distribution of bounding-box area, and the number of background-only images. Object size is expressed as bounding-box area relative to image area:

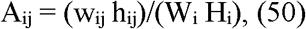

where wᵢⱼ and hᵢⱼ are the width and height of the j-th ground-truth box in image Iᵢ and Wᵢ and Hᵢ the image dimensions. The normalized area satisfies 0 < Aᵢⱼ ≤ 1. Reporting normalized rather than absolute area prevents image resolution from confounding comparisons across differently sized images.

A separate reliability experiment is reported in the Results. Its exported prediction tables contain 122,723 validation detections from 2,147 images and 122,720 test detections from 2,148 images, and the source paths identify a saved 70:15:15 split. These detection-bearing image counts do not establish the full cohort size, because images without candidate detections may be absent from the tables. The reliability results are therefore not pooled with the 8,000-image experiment of Table 1, and no cross-split performance improvement is inferred. The exact checkpoint identity, complete split manifests, and candidate-generation settings accompany the reproducibility record.

### Compared methods

Table 2 lists the intended detector comparators. The completed detector results and the separate reliability component comparisons are reported separately; architecture-baseline comparisons are not inferred from the planned configurations.

**Table 2.**
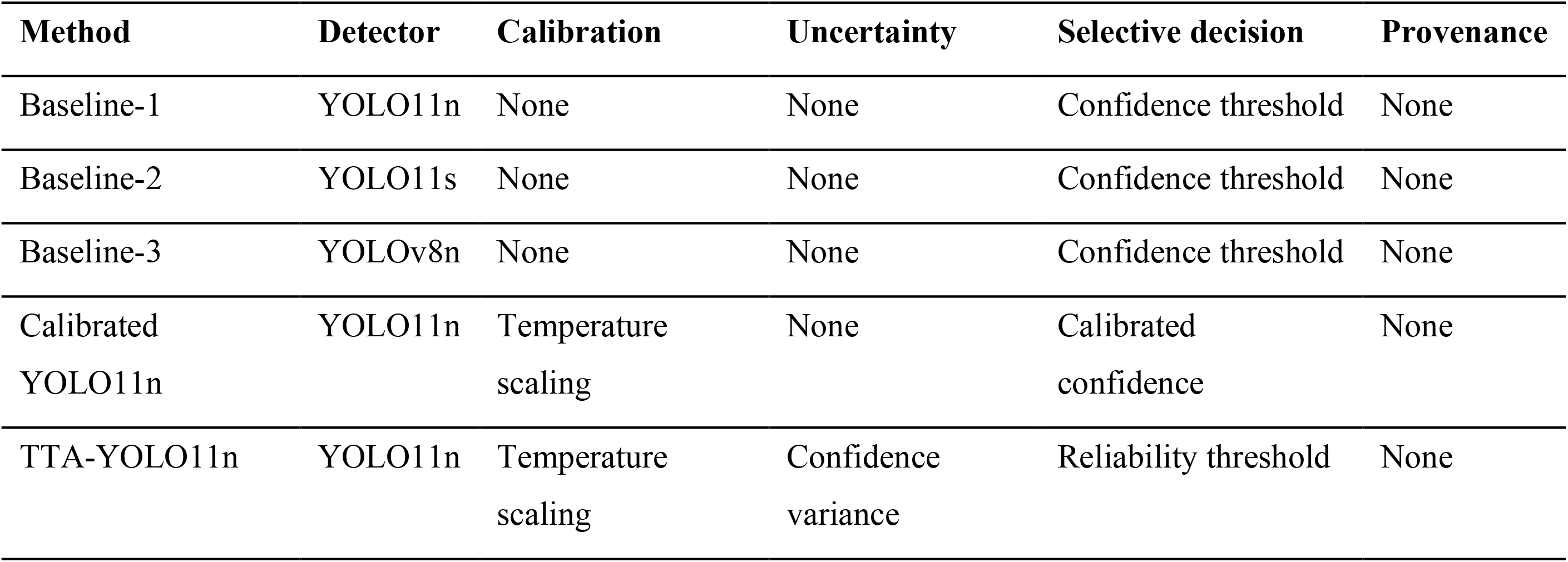
Planned comparator configurations.

| **Method** | **Detector** | **Calibration** | **Uncertainty** | **Selective decision** | **Provenance** |
|---|---|---|---|---|---|
| Baseline-1 | YOLO11n | None | None | Confidence threshold | None |
| Baseline-2 | YOLO11s | None | None | Confidence threshold | None |
| Baseline-3 | YOLOv8n | None | None | Confidence threshold | None |
| Calibrated YOLO11n | YOLO11n | Temperature scaling | None | Calibrated confidence | None |
| TTA-YOLO11n | YOLO11n | Temperature scaling | Confidence variance | Reliability threshold | None |

The contrast between the YOLO11n baseline and WildTrust-UAV isolates the effect of the reliability and provenance modules at fixed detector capacity. YOLO11s tests whether added capacity alone gives a similar operational benefit, and YOLOv8n is an older, lighter comparator. All comparator experiments should use recorded image dimensions, training budgets, stopping criteria, augmentation, and split manifests. Architecture-level comparator measurements are not included in the present results. The reliability component comparisons use the same exported test detections and so required no additional detector training.

### Training configuration

The main detector is trained for up to 100 epochs at 640 × 640 pixels, with early stopping on the validation fitness metric and a patience of 20 epochs. Batch size is set adaptively from GPU memory or fixed at the largest feasible value for all compared models.

The detector experiment reports two completed runs, with seeds 17 and 29; the separate reliability export is not treated as a further repeated-seed comparison:

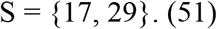

The seed controls initialization, data-loader ordering, sampling, and stochastic augmentation. Deterministic execution is enabled wherever the framework and hardware libraries support it. Because two seed runs were completed, their held-out results are reported individually rather than as a three-seed mean and standard deviation. Table 3 summarizes the configuration.

**Table 3.** Training and inference configuration.

| **Parameter** | **Setting** |
|---|---|
| Input resolution | 640 × 640 |
| Maximum epochs | 100 |
| Early-stopping patience | 20 epochs |
| Pretrained initialization | COCO-pretrained weights |
| Random seeds | 17 and 29 |
| IoU thresholds for evaluation | 0.50 to 0.95 |
| Initial inference confidence | 0.001 for metric construction |
| NMS IoU threshold | 0.70 |
| Optimizer | Ultralytics automatic selection |
| Mixed precision | Enabled when GPU-supported |
| Stochastic inference views | Identity plus four transformations |
| Calibration source | Validation partition only |
| Final evaluation source | Test partition only |

The experimental pipeline records software-library versions, GPU model, CUDA version, operating environment, elapsed training time, peak GPU memory, and the cryptographic hash of the final checkpoint for every seed run, so that the evaluated model can be matched to the reported checkpoint.

### Stochastic inference configuration

Each validation and test image is evaluated with five label-preserving stochastic views:

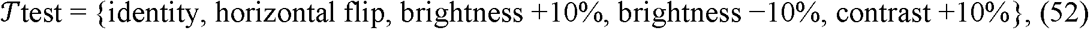

so T = 5. Boxes from each transformed view are mapped back to the original image coordinates before matching, and candidates are matched by class agreement and the IoU threshold

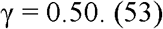

For each reference detection, the analysis records original confidence, correctness, confidence-score variance across matched views, normalized-corner localization variance, matched-view count, detection instability, normalized component scores, composite uncertainty, calibrated confidence, and reliability. Correctness uses confidence-ordered, class-consistent, one-to-one matching to ground truth at IoU 0.50. A reference detection may independently select its highest-IoU same-class candidate in each transformed view.

The composite weights (α, β, δ) are selected on the validation partition by a constrained grid search:

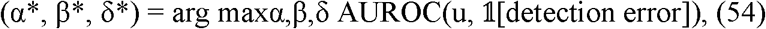

subject to

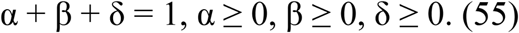

After selection the weights are frozen and the test partition is evaluated once.

### Detection performance measures

Precision and recall are Precision =

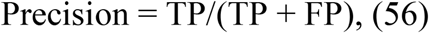

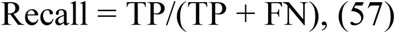

where TP, FP, and FN are the numbers of true positives, false positives, and missed ground-truth instances. Mean average precision at IoU 0.50 is

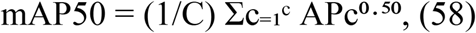

where APc⁰·⁵⁰ is the average precision of class c at IoU 0.50. The stricter COCO-style metric averages over ten IoU thresholds from 0.50 to 0.95 in steps of 0.05:

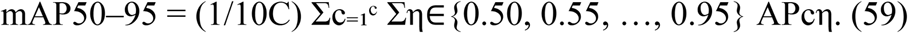

The metrics in Eqs (56) to (59) are reported as macro averages and per class, because aggregate performance can hide poor detection of rare species. Average precision is also stratified by the normalized object area of Eq (50) to show whether gains are concentrated among small targets.

### Calibration evaluation

A detection receives correctness label yₖ = 1 when it matches an unused ground-truth object of the same class with IoU ≥ 0.50, and yₖ = 0 otherwise:

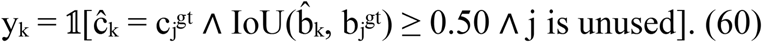

Matching is greedy in descending confidence order; once a ground-truth object is matched it is marked as used, so no two predictions are credited for the same animal. Temperature scaling is fitted on validation detections only and applied unchanged to test detections. Calibration is assessed with the Brier score, NLL, ECE with 10 equal-width bins, maximum calibration error (MCE), and reliability diagrams. With B = 10,

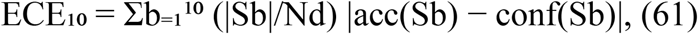

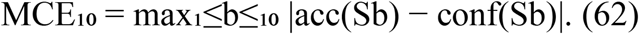

Pre- and post-calibration values are both reported. A fall in ECE₁₀, MCE₁₀, Brier score, and NLL with no loss in mAP supports the calibration stage. mAP is not expected to change, since temperature scaling preserves score ranking; if threshold-dependent metrics change, they are reported separately with thresholds frozen on the validation partition.

### Uncertainty evaluation

Uncertainty quality is evaluated independently of raw accuracy. The primary endpoint is the ability of composite uncertainty uₖ to separate erroneous from correct detections, with error label

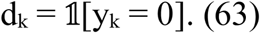

We report the AUROC and area under the precision-recall curve (AUPRC) for identifying erroneous detections, the Spearman correlation between uncertainty and localization error, mean uncertainty for correct and incorrect detections, and uncertainty stratified by normalized object size and by species.

For a prediction matched to a ground-truth box, localization error is

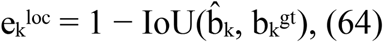

and the expected association is

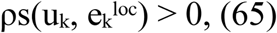

where ρs is Spearman’s rank correlation. A positive value means higher uncertainty goes with worse localization. Correlation alone does not establish probabilistic calibration, so it is read alongside the calibration and selective-risk results. Unmatched predictions have no unique ground-truth box and are excluded from the correlation analysis, but they remain in the error-discrimination analysis of Eq (63).

### Selective-prediction evaluation

Detections are ranked by the reliability score

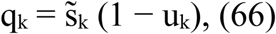

and evaluated at target coverage levels

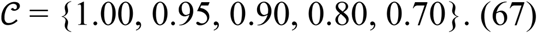

At each level c, detections are sorted by qₖ and the least reliable fraction (1 − c) is deferred. The threshold for coverage c is the validation quantile

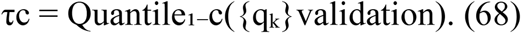

Validation quantiles are applied unchanged to test detections. The reported outputs are target coverage, achieved test coverage, selective risk, and the numbers of accepted and deferred detections. The present export does not provide selective recall or accepted-detection mAP. At nominal full coverage the validation-minimum threshold excludes one test detection, so achieved coverage is 0.999992 rather than 1.0. Threshold ties and distribution differences are handled by reporting achieved coverage explicitly.

### Provenance threat model and attack simulation protocol

To test whether the Merkle-batched, Ed25519-signed provenance mechanism detects the tampering it is designed for, we implemented the pipeline of Eqs (41) to (49): canonical record construction, SHA-256 leaf and Merkle hashing, Ed25519 batch signing, and the joint leaf-recomputation, Merkle-inclusion, and signature verifier. The pipeline was run with the trained YOLO11n seed-17 checkpoint on 32 real WAID test images, so the image, model, configuration, and prediction commitments come from real detector inference. Because WAID has no GPS metadata, the location-policy coordinates are deterministic synthetic values and are labelled as such. The resulting ledger is a local, permissioned, hash-linked prototype; a Hyperledger Fabric payload was exported, but no distributed-consensus transaction is claimed.

Nine adversarial scenarios were simulated on copies of the 32-record real-inference batch:

1. Image substitution: the source-image hash hI in an accepted record is replaced with the hash of a different image, simulating substitution of the underlying photograph after inference.
2. Model switching: the model hash hM and version vM are replaced, simulating a detection falsely attributed to a different or unauthorized model.
3. Prediction tampering: the predicted class and confidence are altered after the fact, simulating manual relabelling or confidence inflation.
4. Configuration modification: the configuration hash hC is replaced, simulating an undisclosed change to preprocessing, tiling, or thresholding.
5. Location-commitment tampering: the salted geographic commitment hgeo is replaced, simulating falsification of the protected coordinate commitment.
6. Timestamp backdating: the inference timestamp is shifted by 24 hours, simulating misrepresentation of when an observation was made.
7. Signature forgery: a fabricated batch root is signed with an unauthorized private key, simulating an unauthenticated device or actor.
8. Merkle-proof forgery: a record never included in a batch is paired with a random authentication path and presented as included under the true, signed root.
9. Replay of a prior valid record: a genuine, previously accepted record, proof, and signature from an earlier batch are resubmitted unchanged, simulating reuse of stale evidence as a new observation; detection relies on the nonce-freshness check rather than a hash or signature mismatch.

Each scenario ran for 500 independent trials, with the attack applied to a selected source record. A further 500-trial control with no tampering measured the false-rejection rate on genuine records. Detection counts as successful when full verification (leaf-hash recomputation with Merkle-inclusion and signature checks, plus freshness bookkeeping for the replay scenario) rejects the submission. Per-record verification latency was recorded for every trial. Merkle-batch construction and Ed25519 signing time were profiled separately at batch sizes of 16, 64, 256, and 1,024 records (100 repeats each). The 16-record batch uses unique real source records; larger batches reuse records derived from the 32 real ones solely to measure scaling.

## Results

### Held-out test detection performance

The YOLO11n detector was trained on the 4,000-image training partition and monitored on the 2,000-image validation partition. Two seed runs were completed: seed 17 stopped after 50 saved epochs, and seed 29 completed the 100-epoch schedule after a CPU auto-resume. Both best checkpoints were evaluated on the same held-out 2,000-image test partition, which contains 33,119 annotated instances. Neither temperature scaling nor the selective gateway was applied to the detection metrics reported here. Table 4 gives the results, computed as in Eqs (56) to (59).

**Table 4.** Held-out test detection performance of the WildTrust-UAV YOLO11n detector for two completed seeds.

| **Metric** | **Seed 17** | **Seed 29** |
|---|---|---|
| Precision | 0.896 | 0.897 |
| Recall | 0.832 | 0.840 |
| mAP@0.50 | 0.901 | 0.903 |
| mAP@0.50:0.95 | 0.527 | 0.527 |

### Scope of the reported results

Table 4 reports the detector experiment and does not establish improvement over the architecture comparators of Table 2. The uncertainty, calibration, selective-prediction, and component-comparison results below come from a separate reliability experiment with a different test image count, and the two experiments are interpreted separately. Two detector runs with different training durations do not support formal claims about seed robustness or statistical significance.

### End-to-end provenance attack detection

Table 5 reports the nine attack scenarios, each run for 500 trials on records derived from the 32-image real-inference batch, together with the 500-trial genuine control.

**Table 5.** Tamper-detection results for the Merkle/Ed25519 provenance mechanism integrated with real detector inference (500 trials per scenario).

| **Attack scenario** | **Trials** | **Detected** | **Detection rate** | **Mean verification latency (ms)** | **Detection mechanism** |
|---|---|---|---|---|---|
| Image substitution (hI swapped) | 500 | 500 | 100.00% | 0.066 | Leaf-hash mismatch (Eq 49) |
| Model switching (hM swapped) | 500 | 500 | 100.00% | 0.105 | Leaf-hash mismatch (Eq 49) |
| Prediction tampering (class/confidence altered) | 500 | 500 | 100.00% | 0.066 | Leaf-hash mismatch (Eq 49) |
| Configuration modification (hC swapped) | 500 | 500 | 100.00% | 0.061 | Leaf-hash mismatch (Eq 49) |
| Location-commitment tampering (hgeo swapped) | 500 | 500 | 100.00% | 0.059 | Leaf-hash mismatch (Eq 49) |
| Timestamp backdating | 500 | 500 | 100.00% | 0.062 | Leaf-hash mismatch (Eq 49) |
| Signature forgery (unauthorized key) | 500 | 500 | 100.00% | 0.182 | Signature failure (Eq 47) |
| Merkle-proof forgery (unincluded record) | 500 | 500 | 100.00% | 0.074 | Inclusion-proof failure (Eq 48) |
| Replay of a prior valid record | 500 | 500 | 100.00% | 0.242 | Nonce-reuse check |
| Control: genuine, untampered records | 500 | 0 rejected | 0.00% (false-rejection rate) | 0.263 | None |

Every attack was detected in all 500 trials (4,500 of 4,500), and the genuine control produced no false rejections in 500 trials.

### Provenance computational overhead

Table 6 reports mean Merkle-batch construction time, Ed25519 signing time, and combined per-record cryptographic cost at four batch sizes (100 repeats each), addressing the provenance part of RQ7. For batch sizes above 32, source records were derived deterministically from the 32 real-inference records for scaling only.

**Table 6.** Provenance batch-construction and signing overhead by batch size (mean of 100 repeats).

| **Batch size (records)** | **Merkle-build time (ms)** | **Ed25519 signing time (ms)** | **Per-record cost (ms)** |
|---|---|---|---|
| 16 | 0.034 | 0.067 | 0.0063 |
| 64 | 0.123 | 0.082 | 0.0032 |
| 256 | 0.756 | 0.175 | 0.0036 |
| 1,024 | 2.295 | 0.156 | 0.0024 |

Signing time stayed between 0.067 and 0.175 ms across batch sizes because one signature is produced per batch. Merkle construction rose from 0.034 ms at 16 records to 2.295 ms at 1,024 records. As the cost is amortized over the batch, the combined cryptographic cost remained between about 0.0024 and 0.0063 ms per record.

### RQ2: Uncertainty discrimination

In the reliability experiment, the composite uncertainty score separated erroneous from correct test detections with an AUROC of 0.9041 and an AUPRC of 0.9515 (Table 7). The reported Spearman correlation between uncertainty and localization error was 0.6290. The test candidate pool contained 122,720 detections with an error prevalence of 0.7221, so the AUPRC should be read against that prevalence. The correlation value is taken from the summary export rather than recomputed from the prediction tables.

**Table 7.**
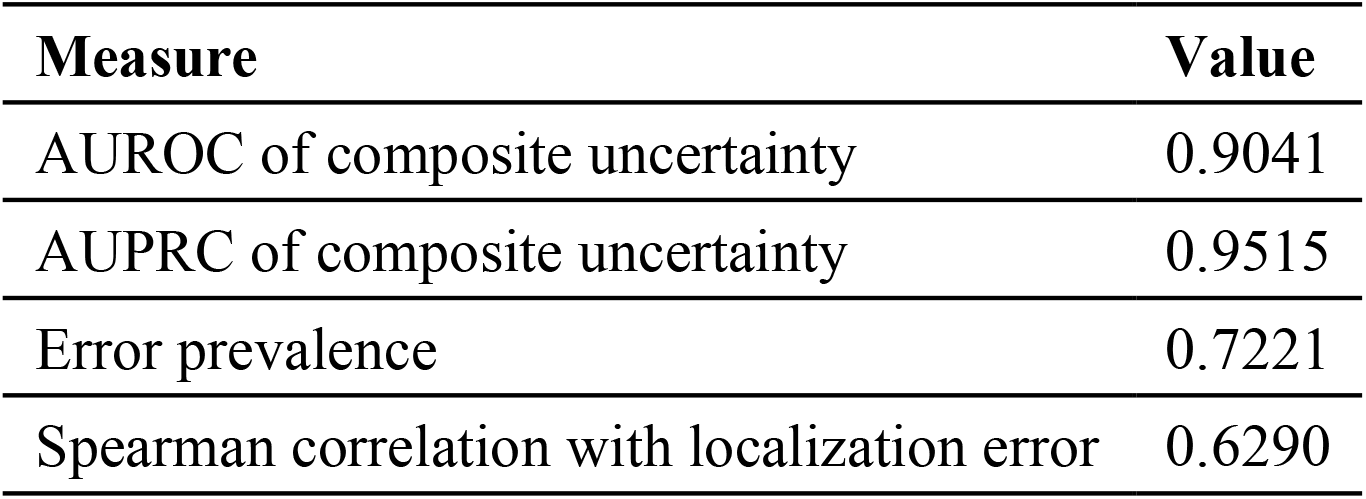
Test error discrimination in the separate reliability experiment.

### RQ3: Confidence calibration

Temperature scaling on validation detections selected T = 0.429168. On test detections, ECE₁₀ fell from 0.071468 to 0.017895, a 74.96% reduction; the Brier score fell from 0.035250 to 0.022230 and NLL from 0.139630 to 0.100497 (Table 8; Fig 3). These measures describe correctness calibration of the retained candidate detections, not of all wildlife observations.

**Fig 3.**
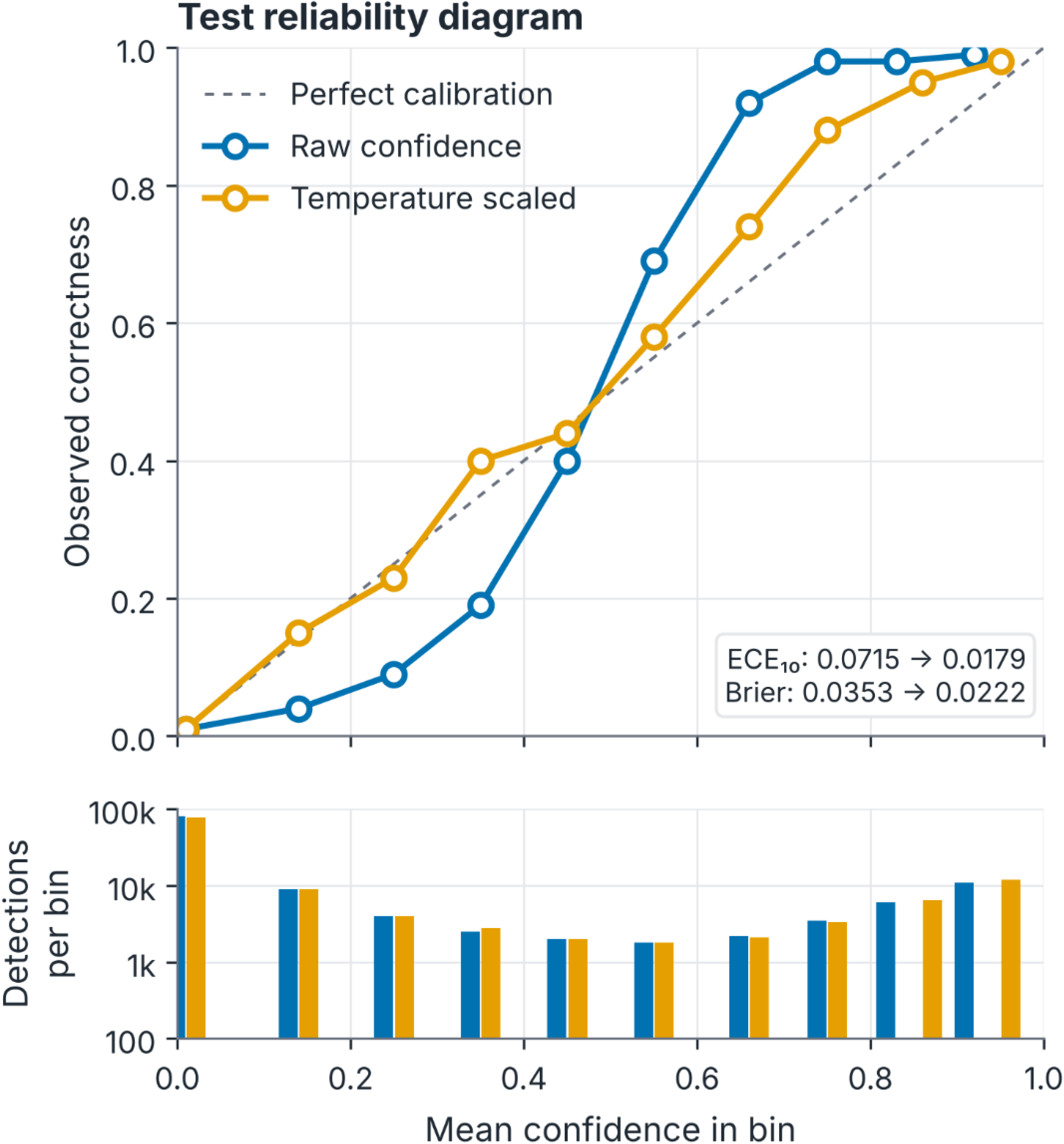
Test reliability diagram for raw and temperature-scaled confidence in the separate reliability experiment. Points are non-empty equal-width confidence bins; the dashed diagonal marks perfect calibration

**Table 8.** Raw and temperature-scaled calibration metrics.

| **Split** | **Stage** | **Brier** | **NLL** | **ECE<sub>10</sub>** | **MCE<sub>10</sub>** |
|---|---|---|---|---|---|
| Validation | Raw | 0.035008 | 0.139321 | 0.071866 | 0.260740 |
| Validation | Temperature scaled | 0.022056 | 0.099608 | 0.017955 | 0.105505 |
| Test | Raw | 0.035250 | 0.139630 | 0.071468 | 0.262696 |
| Test | Temperature scaled | 0.022230 | 0.100497 | 0.017895 | 0.120789 |

### RQ4: Selective prediction

Validation-derived thresholds lowered test selective risk from 0.722056 at near-complete coverage to 0.610172 at a coverage of 0.699356 (Table 9; Fig 4a). The absolute difference is 0.111884, or 11.19 percentage points, with 36,895 detections deferred and 85,825 retained. Residual risk remains high. These risk values are not comparable with the precision operating point in Table 4.

**Fig 4.**
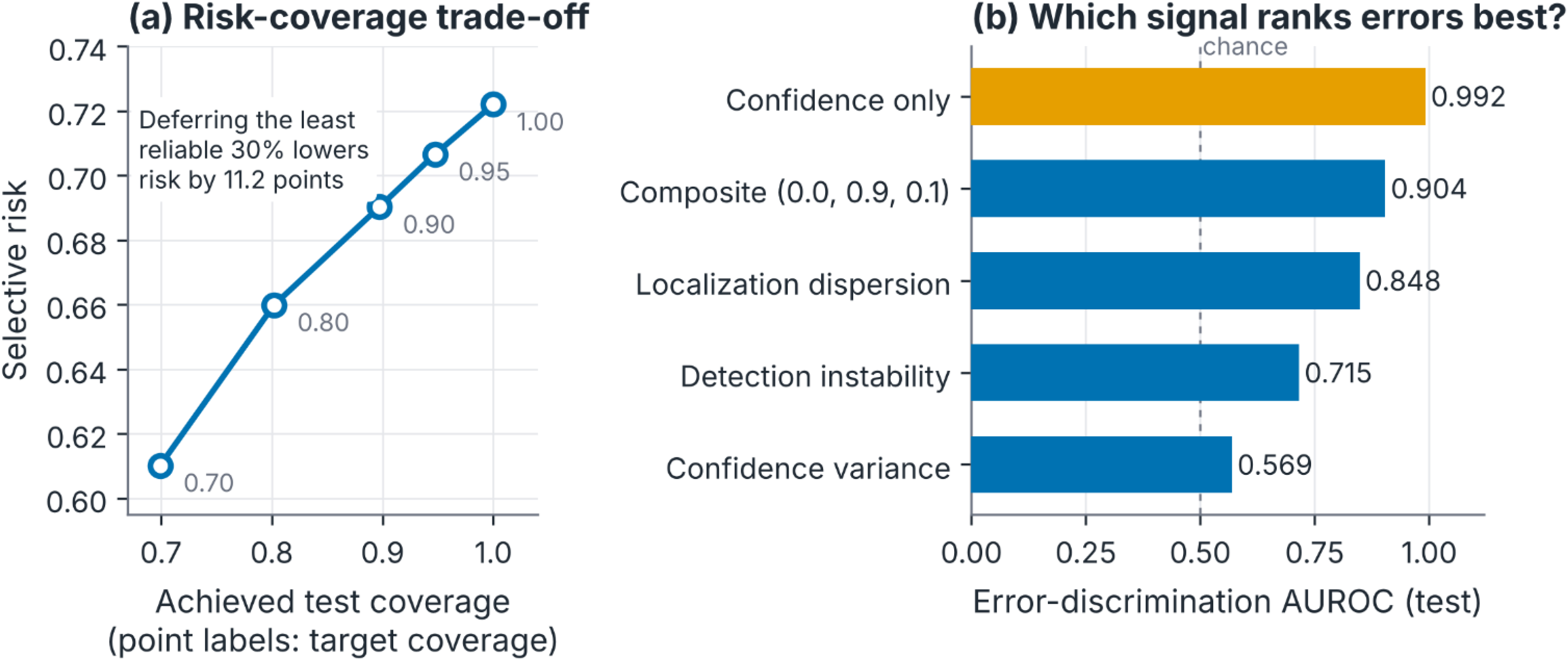
Selective prediction and uncertainty component comparison in the separate reliability experiment. (a) Selective risk at achieved test coverage using validation-derived thresholds. (b) Error-discrimination AUROC for confidence alone and for each uncertainty component and the composite. Values are point estimates without confidence intervals.

**Table 9.** Selective-prediction trade-off using validation-frozen thresholds.

| **Target coverage** | **Validation threshold** | **Test coverage** | **Selective risk** | **Accepted** | **Deferred** |
|---|---|---|---|---|---|
| 1.00 | 9.227 × 10<sup>−20</sup> | 0.999992 | 0.722056 | 122,719 | 1 |
| 0.95 | 2.518 × 10<sup>−19</sup> | 0.947254 | 0.706607 | 116,247 | 6,473 |
| 0.90 | 1.261 × 10<sup>−18</sup> | 0.897164 | 0.690263 | 110,100 | 12,620 |
| 0.80 | 1.087 × 10<sup>−8</sup> | 0.802094 | 0.659901 | 98,433 | 24,287 |
| 0.70 | 3.149 × 10<sup>−7</sup> | 0.699356 | 0.610172 | 85,825 | 36,895 |

### RQ5: Uncertainty component comparison

Validation selected α = 0.0, β = 0.9, and δ = 0.1. On the test set, localization dispersion alone reached an AUROC of 0.8483, instability alone 0.7154, and confidence variance alone 0.5689; the composite reached 0.9041. Confidence-only error ranking, however, reached an AUROC of 0.9917 and an AUPRC of 0.9964, exceeding the composite (Table 10; Fig 4b). The experiment therefore supports the narrower claim that localization dispersion and instability together rank errors better than either alone, and shows that confidence alone remains the strongest single ranking signal in this setting.

**Table 10.** Test error discrimination for confidence and uncertainty components.

| **Variant** | **AUROC (error)** | **AUPRC (error)** |
|---|---|---|
| Confidence only | 0.991684 | 0.996431 |
| Classification (confidence variance) | 0.568932 | 0.830216 |
| Localization | 0.848301 | 0.932466 |
| Stability | 0.715361 | 0.838223 |
| Composite | 0.904121 | 0.951527 |

WildTrust-UAV examines how an aerial wildlife prediction can be turned into a more reliable and verifiable observation. Its components map onto successive steps of the conservation workflow: detection identifies animals, calibration and uncertainty estimate how far each prediction can be trusted, the selective gateway decides which observations need human validation, and the provenance record documents how each observation was produced. The detection experiment, the provenance experiment, and the separate reliability assessment provide complementary evidence about these functions. The results are discussed below in relation to the research questions.

### RQ1: Detection performance

In the detector experiment, two completed YOLO11n runs reached mAP@0.50 of 0.901 and 0.903 on the 2,000-image test set, indicating consistent detection performance. This matters for the downstream modules, because uncertainty and evidence assessments are only meaningful when applied to credible candidate observations. The stricter mAP@0.50:0.95 of 0.527 shows a sizeable gap between finding an animal and localizing it precisely, which is common when targets occupy little of the frame. WildTrust-UAV uses localization variance as part of its reliability score for this reason, combining spatial consistency with the confidence value. The two runs are reported separately because their training durations differed.

### RQ2: Uncertainty discrimination

The composite uncertainty score produced an AUROC of 0.9041, meaning that wrong candidate detections received higher uncertainty on average than correct ones. This supports its use for prioritizing expert inspection: in surveys with many candidate detections, a ranked review queue is more practical than an unordered one.

The components capture different aspects of prediction behaviour. Confidence variance reflects score instability, localization dispersion reflects boundary instability, and detection instability reflects whether the animal is found at all under transformation. Together they describe how a prediction behaves under mild changes to the image.

Confidence-only ranking achieved a higher AUROC of 0.9917 in this experiment. Confidence therefore remains the strongest basis for error ranking, and the composite adds a description of prediction variability rather than a better ranking. The AUPRC of 0.9515 characterizes error retrieval among candidates with an error prevalence of 0.7221. These results separate two questions that are often conflated: what uncertainty information is worth, and which score ranks errors best.

### RQ3: Calibration

Calibration substantially improved the interpretability of detector confidence. Temperature scaling fitted on validation samples reduced test ECE from 0.0715 to 0.0179, about 75%, and the Brier score from 0.0353 to 0.0222. NLL and MCE improved as well. The change matters because confidence values can decide whether an observation is accepted, referred, or deferred. Better agreement between confidence and correctness gives a sounder basis for setting those thresholds. Because the temperature was fitted on validation predictions and evaluated on test predictions, the improvement extends beyond the data used to fit it. For the evaluated candidate pool, calibration improved the informativeness of the confidence scores.

### RQ4: Selective prediction

The selective prediction results quantify the trade-off between automation coverage and the error rate of accepted predictions. Risk fell from 0.7221 to 0.6102, a reduction of 11.19 percentage points, at 69.94% coverage.

This gives conservation teams a quantitative basis for deciding how much to defer to manual review. Different survey goals justify different operating points, depending on whether broader coverage or more selective acceptance is preferred. The findings describe an empirical trade-off rather than a universal threshold: the operating points were evaluated on a candidate pool that includes many low-confidence detections, and substantial residual error remains. The practical value lies in making the consequences of deferral measurable.

### RQ5: Component contribution

The validation search assigned 0.9 of the composite weight to localization dispersion, 0.1 to detection instability, and none to confidence variance. Localization consistency is therefore the main contributor in the chosen combination.

This pattern fits aerial wildlife imagery, where variation in predicted boundaries can signal ambiguity in small or partly occluded targets, and detection instability adds information about consistency across views. Their combination reached an AUROC of 0.9041, above localization alone (0.8483) and instability alone (0.7154), so the two signals complement each other. Confidence ranking performed better still regardless of the components used. The comparison therefore explains how the uncertainty module behaves without showing that any combination is always superior, and it suggests that a simplified module built on the effective signals would lose little.

### RQ6: Spatial disclosure and integrity

Observations serve several stakeholders who need different degrees of location precision. Role-based disclosure in WildTrust-UAV makes the precision of released coordinates depend on the recipient. The location commitment works alongside the disclosure mechanism and makes any change to committed location information detectable. In the tests, location-commitment tampering was rejected in every trial, showing that spatial metadata can form part of the verifiable evidence record. Disclosure control and commitment verification thus play complementary roles: one governs how much detail is released, the other confirms what was committed. Synthetic coordinates were used because WAID has no field GPS metadata, so the experiment demonstrates how the disclosure and integrity mechanisms operate; a field test remains separate work.

### RQ7: Overhead and verification

All 4,500 tampered submissions across nine scenarios were rejected and all 500 valid records were accepted, so the verifier distinguished the tested alterations from genuine records. Binding the image, model, inference setup, predictions, and metadata makes each evidence record traceable to its computational source. Merkle aggregation makes this cheap, since many records share one signed batch commitment. Combined batch-construction and signing time was 0.101 ms for 16 records and 2.451 ms for 1,024 records in the prototype, so commitment and signing add little computational cost at these batch sizes.

The timing covers local record construction and signing, not full inference or distributed storage. The results show that verifiable provenance can be added to a wildlife analysis pipeline. Provenance authenticates the inference history; detection and reliability evaluation address whether the prediction is accurate.

### Limitations

Several limits on the present evidence should be stated plainly. The detector experiment and the reliability experiment use different data splits and image counts and are not pooled. Only two detector seeds were completed, with different training durations, so no formal seed-robustness or significance claims are made, and the planned architecture comparators of Table 2 have not been measured. The reliability results are point estimates without confidence intervals. The selective-prediction export does not include selective recall or accepted-detection mAP, and residual risk after deferral remains high because the candidate pool includes many low-confidence detections. Location policies were exercised with synthetic coordinates, and the ledger is a local hash-linked prototype rather than a deployed distributed ledger. Field validation under real survey conditions remains to be done.

## Conclusions

Reliable wildlife monitoring requires more than detecting animals in UAV imagery. A system must say how much each prediction can be trusted, refer uncertain detections to experts, release sensitive locations appropriately, and keep each inference traceable to its source. WildTrust-UAV addresses these needs in one framework that combines wildlife detection, calibrated confidence, uncertainty-driven referral, role-based spatial disclosure, and cryptographically verifiable provenance. Its central idea is the link between prediction quality and evidence credibility: an observation can be judged for reliability while keeping a verifiable connection to the image, model, configuration, and metadata that produced it.

The experiments support this design through complementary assessments. The detector reached mAP@0.50 of 0.901 to 0.903, giving a sound detection baseline. In the separate reliability experiment, temperature scaling reduced test ECE by about 75% (0.0715 to 0.0179) and improved the Brier score from 0.0353 to 0.0222. Composite uncertainty detected errors with an AUROC of 0.9041, and selective referral lowered risk among accepted candidates by 11.19 percentage points at 69.94% coverage, although confidence alone ranked errors better than the composite. The provenance verifier rejected all 4,500 tampered submissions and accepted all 500 genuine controls, with batch construction and signing taking between 0.101 and 2.451 ms. Role-based disclosure extended the evidence-handling framework to sensitive location data.

The current evidence comes from benchmark experiments and prototype provenance implementation with synthetic coordinates for the spatial tests. Field experiments are needed to evaluate the framework under operational ecological conditions.

## Acknowledgments

The authors thank the College of Information Sciences and Technology, Pennsylvania State University, and Graphic Era University, India, for the institutional support that made this work possible.

## Data Availability Statement

The public 8,000-image WAID-derived subset used in the experiments is based on the WAID dataset at the link - https://github.com/xiaohuicui/WAID

## References

1. Colomina I, Molina P. Unmanned aerial systems for photogrammetry and remote sensing: A review. ISPRS J Photogramm Remote Sens. 2014;92: 79–97. doi: 10.1016/j.isprsjprs.2014.02.013

2. Hodgson JC, Mott R, Baylis SM, Pham TT, Wotherspoon S, Kilpatrick AD, et al. Drones count wildlife more accurately and precisely than humans. Methods Ecol Evol. 2018;9: 1160–1167. doi: 10.1111/2041-210X.12974

3. Kellenberger B, Marcos D, Tuia D. Detecting mammals in UAV images: Best practices to address a substantially imbalanced dataset with deep learning. Remote Sens Environ. 2018;216: 139–153. doi: 10.1016/j.rse.2018.06.028

4. Bondi E, Jain R, Aggrawal P, Anand S, Hannaford R, Kapoor A, et al. BIRDSAI: A dataset for detection and tracking in aerial thermal infrared videos. In: Proceedings of the IEEE Winter Conference on Applications of Computer Vision (WACV); 2020 Mar 1–5; Snowmass Village, CO, USA. Piscataway: IEEE; 2020. pp. 1747–1756. doi: 10.1109/WACV45572.2020.9093284

5. Mou C, Liu T, Zhu C, Cui X. WAID: A large-scale dataset for wildlife detection with drones. Appl Sci. 2023;13: 10397. doi: 10.3390/app131810397

6. Kumar S, Zhang B, Gudavalli C, Levenson C, Hughey L, Stabach JA, et al. WildlifeMapper: Aerial image analysis for multi-species detection and identification. In: Proceedings of the IEEE/CVF Conference on Computer Vision and Pattern Recognition (CVPR); 2024 Jun 16–22; Seattle, WA, USA. Piscataway: IEEE; 2024. pp. 12594–12604. doi: 10.1109/CVPR52733.2024.01197

7. Guo C, Pleiss G, Sun Y, Weinberger KQ. On calibration of modern neural networks. In: Proceedings of the 34th International Conference on Machine Learning (ICML); 2017 Aug 6–11; Sydney, Australia. PMLR; 2017. Vol. 70, pp. 1321–1330.

8. Geifman Y, El-Yaniv R. Selective classification for deep neural networks. In: Advances in Neural Information Processing Systems 30 (NeurIPS 2017); 2017 Dec 4–9; Long Beach, CA, USA. 2017.

9. Xu D, Ren N, Zhu C. Integrity authentication based on blockchain and perceptual hash for remote-sensing imagery. Remote Sens. 2023;15: 4860. doi: 10.3390/rs15194860

10. Benet J. IPFS: Content addressed, versioned, P2P file system. arXiv:1407.3561 [Preprint]. 2014. Available from: https://arxiv.org/abs/1407.3561

## References

1. Merkle RC. A digital signature based on a conventional encryption function. In: Pomerance C, editor. Advances in Cryptology, CRYPTO ’87. Berlin: Springer; 1988. pp. 369–378. doi: 10.1007/3-540-48184-2_32

2. Josefsson S, Liusvaara I. Edwards-Curve Digital Signature Algorithm (EdDSA). RFC 8032. Internet Research Task Force; 2017. doi: 10.17487/RFC8032

3. Tuia D, Kellenberger B, Beery S, Costelloe BR, Zuffi S, Risse B, et al. Perspectives in machine learning for wildlife conservation. Nat Commun. 2022;13: 792. doi: 10.1038/s41467-022-27980-y

4. Eikelboom JAJ, Wind J, van de Ven E, Kenana LM, Schroder B, de Knegt HJ, et al. Improving the precision and accuracy of animal population estimates with aerial image object detection. Methods Ecol Evol. 2019;10: 1875–1887. doi: 10.1111/2041-210X.13277

5. Kellenberger B, Marcos D, Lobry S, Tuia D. Half a percent of labels is enough: Efficient animal detection in UAV imagery using deep CNNs and active learning. IEEE Trans Geosci Remote Sens. 2019;57: 9524–9533. doi: 10.1109/TGRS.2019.2927393

6. Kellenberger B, Tuia D, Morris D. AIDE: Accelerating image-based ecological surveys with interactive machine learning. Methods Ecol Evol. 2020;11: 1716–1727. doi: 10.1111/2041-210X.13489

7. Lakshminarayanan B, Pritzel A, Blundell C. Simple and scalable predictive uncertainty estimation using deep ensembles. In: Advances in Neural Information Processing Systems 30 (NeurIPS 2017); 2017 Dec 4–9; Long Beach, CA, USA. 2017. pp. 6402–6413.

8. Geifman Y, El-Yaniv R. SelectiveNet: A deep neural network with an integrated reject option. In: Proceedings of the 36th International Conference on Machine Learning (ICML); 2019 Jun 9–15; Long Beach, CA, USA. PMLR; 2019. Vol. 97, pp. 2151–2159.

9. Chapman AD, Grafton O. Guide to best practices for generalising sensitive species occurrence data. Copenhagen: Global Biodiversity Information Facility; 2008.

10. De Campos MGS, Chanel CPC, Chauffaut C, Lacan J. Towards a blockchain-based multi-UAV surveillance system. Front Robot AI. 2021;8: 557692. doi: 10.3389/frobt.2021.557692

11. Liang X, Shetty S, Tosh D, Kamhoua C, Kwiat K, Njilla L. ProvChain: A blockchain-based data provenance architecture in cloud environment with enhanced privacy and availability. Future Gener Comput Syst. 2017;67: 270–281. doi: 10.1016/j.future.2016.10.003

12. Moreau L, Missier P, editors. PROV-DM: The PROV data model. W3C Recommendation. Cambridge (MA): World Wide Web Consortium; 2013.

